# C2C12 muscle myotubes, but not kidney proximal tubule HK-2 cells, elevate erythritol synthesis in response to oxidative stress

**DOI:** 10.1101/2023.07.18.549518

**Authors:** Semira R. Ortiz, Martha S. Field

**Author notes:** **Corresponding author:** Martha S. Field, 113 Savage Hall, Division of Nutritional Sciences, Ithaca, NY 14853, USA. **Author last names:** Ortiz, Field.

## Abstract

**Background:** As a biomarker, elevated serum erythritol predicts type 2 diabetes and cardiovascular disease onset. Erythritol was recently shown to be a product of human glucose metabolism through the pentose phosphate pathway. The regulation of erythritol synthesis from glucose has been explored in cancer cells, but not in non-transformed cells.

**Objective:** The kidneys and skeletal muscle have increased erythritol content in response to dietary sucrose, which suggests that they may significantly contribute to circulating erythritol levels. In the present study, we evaluated if conditions that promote erythritol synthesis in cancer cells are consistent in skeletal muscle and kidney cells.

**Methods:** C2C12 myotubules were used as a model for skeletal muscle and HK-2 human proximal tubule cells were used to model kidney. C2C12 cells were exposed to high-or low-glucose conditions. Both C2C12 and HK-2 cells were exposed to the free radical generator menadione, then intracellular reactive oxygen species (ROS) and erythritol were measured. Intracellular sorbitol levels were also measured because increased polyol flux is also observed after exposure to excess glucose and oxidative stress.

**Results:** Intracellular erythritol was significantly elevated in C2C12 cells following both high glucose and menadione treatment. In contrast, HK-2 cells did not increase erythritol synthesis in response to oxidative stress. Generation of ROS through hydrogen peroxide (H_2_O_2_) exposure elevated sorbitol levels in both C2C12 and HK-2 cells, whereas generation of radicals with menadione treatment did not affect sorbitol production in either cell type.

**Conclusions:** These findings highlight that the factors contributing to elevated erythritol synthesis vary between cell types. More specifically, these studies demonstrate that muscle cells increase erythritol synthesis in response to both high glucose in culture medium and oxidative stress, whereas kidney cells increase erythritol synthesis only in response to high glucose.

## Introduction

Erythritol is synthesized from glucose through the pentose phosphate pathway (PPP) (1–3). The regulation of erythritol synthesis has been characterized in human lung cancer cells, but these mechanisms have not been validated in non-transformed tissues or cells (1,3). Previous work in mice indicates that the liver, kidneys, and skeletal muscle may be major sites of erythritol synthesis in mammals (4). It was also recently found that both mouse kidneys and the skeletal muscle respond to high sucrose intake with elevated erythritol content, whereas the liver maintains a constant erythritol level even after exposure to excess sucrose (5). Sugar-induced erythritol synthesis in muscle is surprising: in healthy adult muscle, there is relatively little PPP activity, and erythritol synthesis from glucose depends on the PPP (6,7). Understanding the role of erythritol production in skeletal muscle is important due to the strong contribution of muscle to glucose disposal/homeostasis. If erythritol is an additional point of disposal for sugar during nutrient excess, skeletal muscle is likely to be a strong contributor to circulating erythritol levels.

While elevated kidney erythritol was expected in response to sucrose intake, several questions remain to be explored. Primarily, it is unclear if kidney erythritol content is reflective of kidney erythritol synthesis, or a result of the urinary clearance of erythritol. There is evidence that proximal tubule cells increase erythritol synthesis in response to high-glucose culture medium, however, further work is required to identify the regulation of erythritol synthesis in kidney cells (3,8).

There may also be interactions between erythritol synthesis and other glucose disposal pathways. One enzyme that catalyzes erythritol synthesis, sorbitol dehydrogenase (SORD), primarily converts glucose-derived sorbitol to fructose through the polyol pathway **(Figure 1)** (9). The polyol pathway becomes overactive in response to hyperglycemia and can contribute to the pathogenesis of diabetic complications (9). Erythritol synthesis through the PPP may be a favorable alternative mechanism of glucose “disposal” due to the rapid excretion of erythritol with limited impact on osmotic stress compared to sorbitol. We have observed that erythritol and sorbitol levels are often inversely associated, however, the relationship between the polyol pathway and erythritol synthesis has not been directly assessed (3).

**Figure 1.**
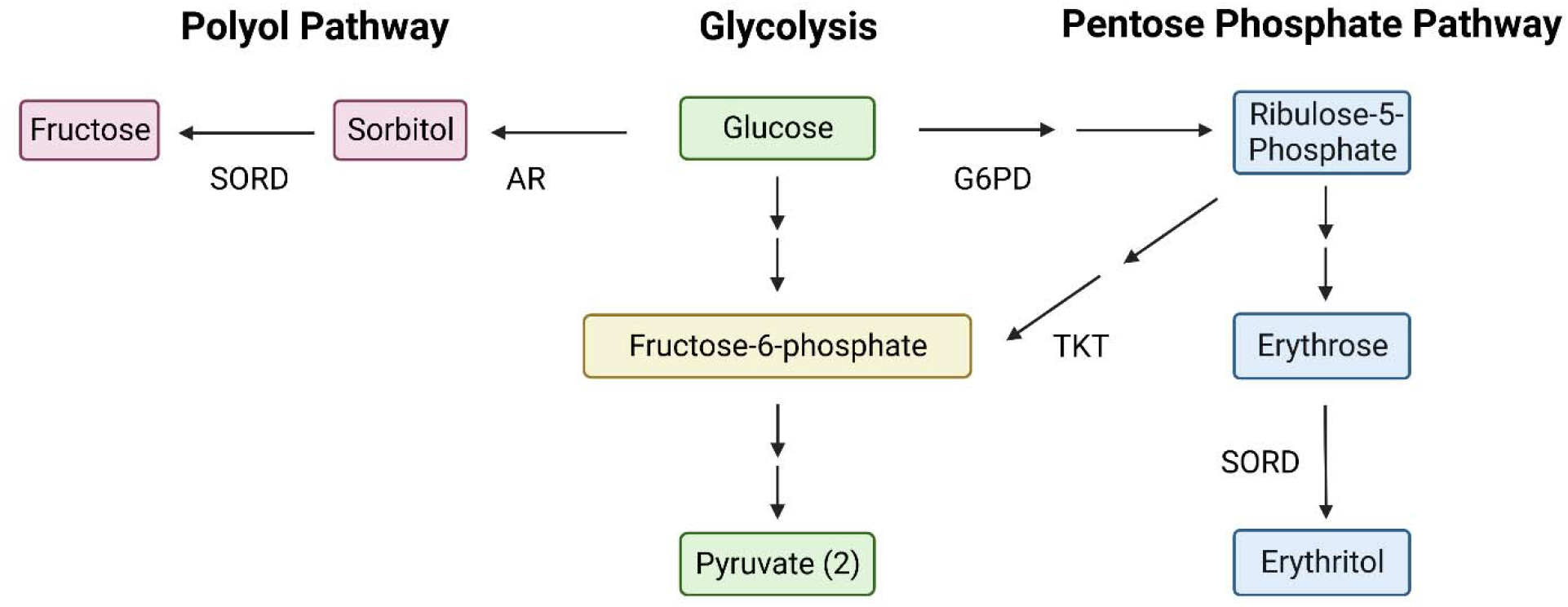
Metabolism of glucose through glycolysis, the polyol pathway, and the pentose phosphate pathway. Sorbitol dehydrogenase (SORD) participates in two pathways that diverge from glycolysis. The canonical function of SORD is to convert sorbitol to fructose in the polyol pathway. In the pentose phosphate pathway, SORD also catalyzes the conversion of erythrose to erythritol. AR: aldose reductase; G6PD: glucose-6-phosphate dehydrogenase; SORD: sorbitol dehydrogenase; TKT: transketolase.

The purpose of this work was to validate mechanisms regulating erythritol synthesis in these physiologically relevant cell types. We also aimed to assess how inhibition of sorbitol synthesis through the polyol pathway impacts erythritol production in these cells. We hypothesized that both skeletal muscle and kidney cells would respond to factors known to promote PPP flux with increased erythritol synthesis, as previously observed in cancer cells. We expected that polyol pathway inhibition would cause an increase in intracellular erythritol.

## Methods

### Cell culture conditions

C2C12 (CRL-1772) cells were obtained from ATCC and maintained in Dulbecco’s Modified Eagle Medium (DMEM) (Gibco) with 1% penicillin/streptomycin, 1% GlutaMAX (Gibco) and 10% FBS (Cytiva). To differentiate into mytotubes, cells were seeded at 80% confluence, then cultured in differentiation media for 4 days. Differentiation media consisted of DMEM, 2% heat-inactivated horse serum (Gibco), 1% GlutaMAX, and 1% penicillin/streptomycin and was changed every 24 hrs. All C2C12 experiments were performed using myotubes that were differentiated for 4 days. HK-2 (CRL-2190) cells were obtained from ATCC and maintained in DMEM/F12 containing 50% DMEM, 50% Ham’s F12 Nutrient Mix (Gibco), 10% FBS, 1% penicillin/streptomycin, 1% GlutaMAX, and 1% ITS-G supplement (Gibco). The final glucose concentration of the combined DMEM/F12 was 16 mM. For metabolite quantification, HK-2 cells were seeded at a density of 1-1.5 × 10^5^ cells per well in 6-well plates.

### Experimental treatments

For experiments using low glucose in C2C12 or HK-2 cells, standard high-glucose media was replaced by low-glucose DMEM (5 mM) or low-glucose DMEM/F12 and cells were incubated for 24 hours before harvesting polar metabolites. To characterized erythritol synthesis during oxidative stress, cells were exposed to menadione sodium bisulfite (Sigma) or hydrogen peroxide (H_2_O_2_) (Thermo Scientific) at the indicated dose for 2 hours, after which polar metabolites were extracted. Cells were incubated with the aldose reductase inhibitor Sorbinil (Sigma) at the indicated dose for 24 hours before harvesting polar metabolites.

### Extraction and measurement of polar metabolites by GC-MS

Polar metabolites were extracted as previously described (3). 10 μM ^13^C_1_-ribitol (Cambridge Isotope Laboratories) was added to methanol as an internal standard during extraction. Dried extracts were derivatized and metabolites (erythritol, ^13^C_1_-ribitol, and sorbitol) were measured by GC-MS as previously described (4). In SIM mode, mass spectra of erythritol (*m/z* 217), ^13^C_1_-ribitol (*m/z* 218), and sorbitol (*m/z* 319) were acquired from 8-9 min, 10-11 min, and 12-13 min, respectively. Metabolite peaks were selected based on the retention time of their respective standards. Relative erythritol and sorbitol were calculated by dividing their absolute intensity by the absolute intensity of ^13^C_1_-ribitol. Relative erythritol and sorbitol were then normalized to total protein, measured by BCA assay.

### Protein quantification

Cell protein pellets were obtained either from the inter phase during metabolite extraction or from replica plated/treated wells and lysed as previously described (3). Protein was quantified using the Pierce BCA Assay (Thermo Scientific) per the manufacturer’s protocol. Briefly, 20 uL of standard or sample were combined with 200 uL of BCA working reagent, incubated for 15 minutes at 65 C, then absorbance at 562 nm was measured.

### Quantification of intracellular reactive oxygen species

Reactive oxygen species (ROS) were measured using the ROS-Glo H_2_O_2_ Assay (Promega) following the manufacturer’s protocol. HK-2 cells were seeded at a density of 5-7.5 × 10^3^ cells in a white-sided 96-well plate and allowed to adhere overnight. Cells were treated with menadione or H_2_O_2_ at the dose indicated for 2-hours with the H_2_O_2_ Substrate Solution. The ROS-Glo Detection Reagent was added and incubated for 20 minutes at room temperature, then luminescence was recorded. Relative luminescence was normalized to total protein content to account for cell number. Protein was extracted using M-PER (Thermo Scientific) following the manufacturer’s protocol and quantified by Pierce BCA as described above.

### Statistical Analysis

Statistical analyses were conducted in GraphPad Prism 9 (GraphPad Software). All data are shown as mean ± SD, and p-values lower than 0.05 were considered statistically significant. Comparisons between two groups were analyzed by unpaired t-test. Comparisons between more than two groups were analyzed by one-way ANOVA followed by Tukey’s multiple comparisons test or two-way ANOVA with Sidak’s or Tukey’s multiple comparisons test.

## Results

### Erythritol is elevated in C2C12 myotubes in response to glucose and menadione

C2C12 cells increased erythritol content by 40 percent when cultured in 25 mM glucose compared to 5 mM glucose (**Figure 2A**, p < 0.01). As expected, intracellular sorbitol was significantly elevated in response to high-glucose media (Figure 2B, p < 0.0001).

**Figure 2.**
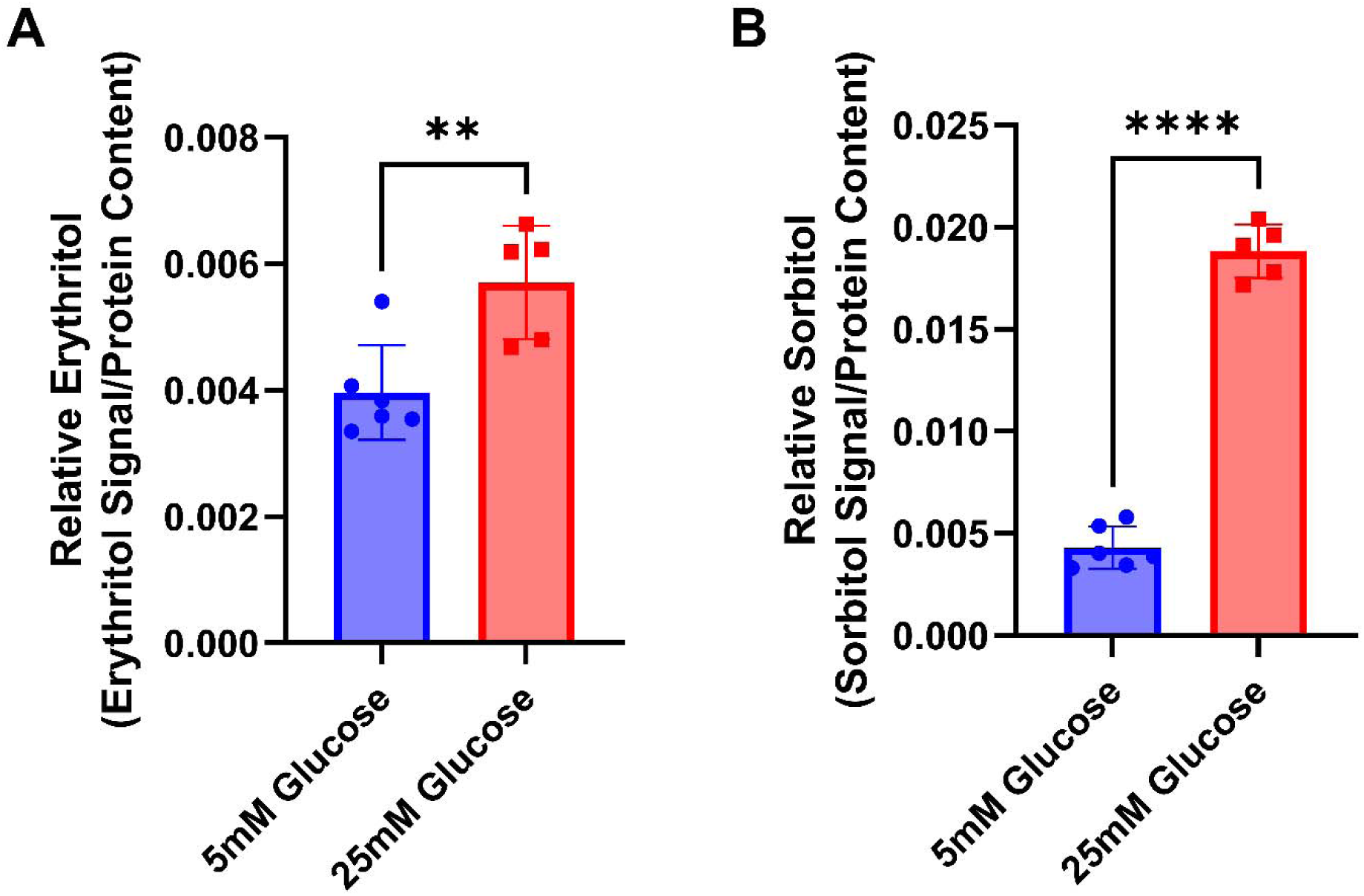
High glucose treatment elevates intracellular erythritol and sorbitol in C2C12 cells. Relative A) erythritol and B) sorbitol in differentiated C2C12 myotubes treated with 5 mM (low) or 25 mM (high) glucose for 24 hours. Relative metabolite values are normalized to internal standard and cell protein content. Data are shown as mean ± SD and were analyzed by unpaired t-test (n=5). **p<0.01, ****p<0.0001.

We utilized two models of elevated reactive oxygen species, direct treatment with H_2_O_2_ and treatment with the free radical generator menadione. Treatment with H_2_O_2_, as expected, resulted in a significant increase in relative ROS (**Figure 3A**, p < 0.0001). Menadione did not significantly elevate relative ROS (Figure 3A) in C2C12 cells. In fact, 20 μM menadione caused a modest, but statistically significant reduction in relative ROS (Figure 3A, p < 0.05). Menadione caused a more than 3-fold increase in intracellular erythritol, but H_2_O_2_ treatment had no impact (Figure 3B, p < 0.0001). Inversely, we observed that H_2_O_2_ treatment significantly elevated sorbitol, whereas menadione treatment had no impact on sorbitol (Figure 3C, p < 0.0001).

**Figure 3.**
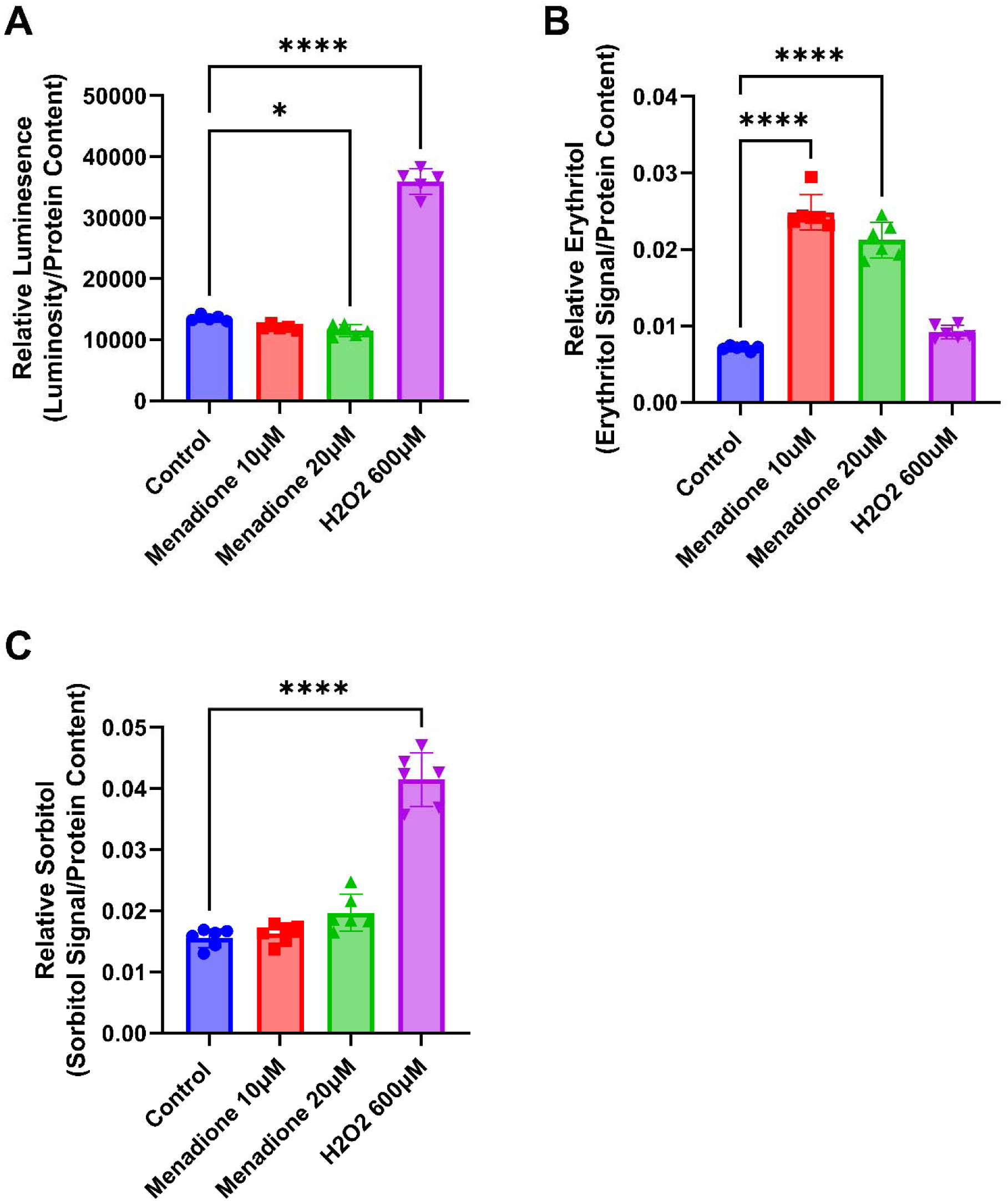
Menadione elevates erythritol content in C2C12 cells. Relative A) ROS, B) erythritol, and C) sorbitol in differentiated C2C12 myotubes treated with menadione or H_2_O_2_ for 2 hrs. Relative metabolite values are normalized to internal standard and all values were normalized to protein content. Data are shown as mean ± SD and were analyzed by one-way ANOVA followed by Tukey’s multiple comparisons test (n=6). *p<0.05, ****p<0.0001. H_2_O_2_: hydrogen peroxide; ROS: reactive oxygen species.

### HK-2 cells do not respond to menadione exposure with elevated erythritol synthesis

Based on findings from A549 and C2C12 cells, we expected that HK-2 cells would also have elevated intracellular erythritol following oxidative stress. We did not, however, observe the same pattern in this cell type. First, we treated cells with 50 uM menadione, which was sufficient to elevated intracellular ROS (**Figure 4A**, p < 0.0001), and 500 μM glutathione ethyl ester (a cell-permeable antioxidant), which was sufficient to reduce intracellular ROS in the presence and absence of menadione (Figure 4A, p < 0.0001). There was no difference in erythritol levels between control and menadione-treated cells (Figure 4B). Glutathione treatment did not significantly impact intracellular erythritol (Figure 4B).

**Figure 4.**
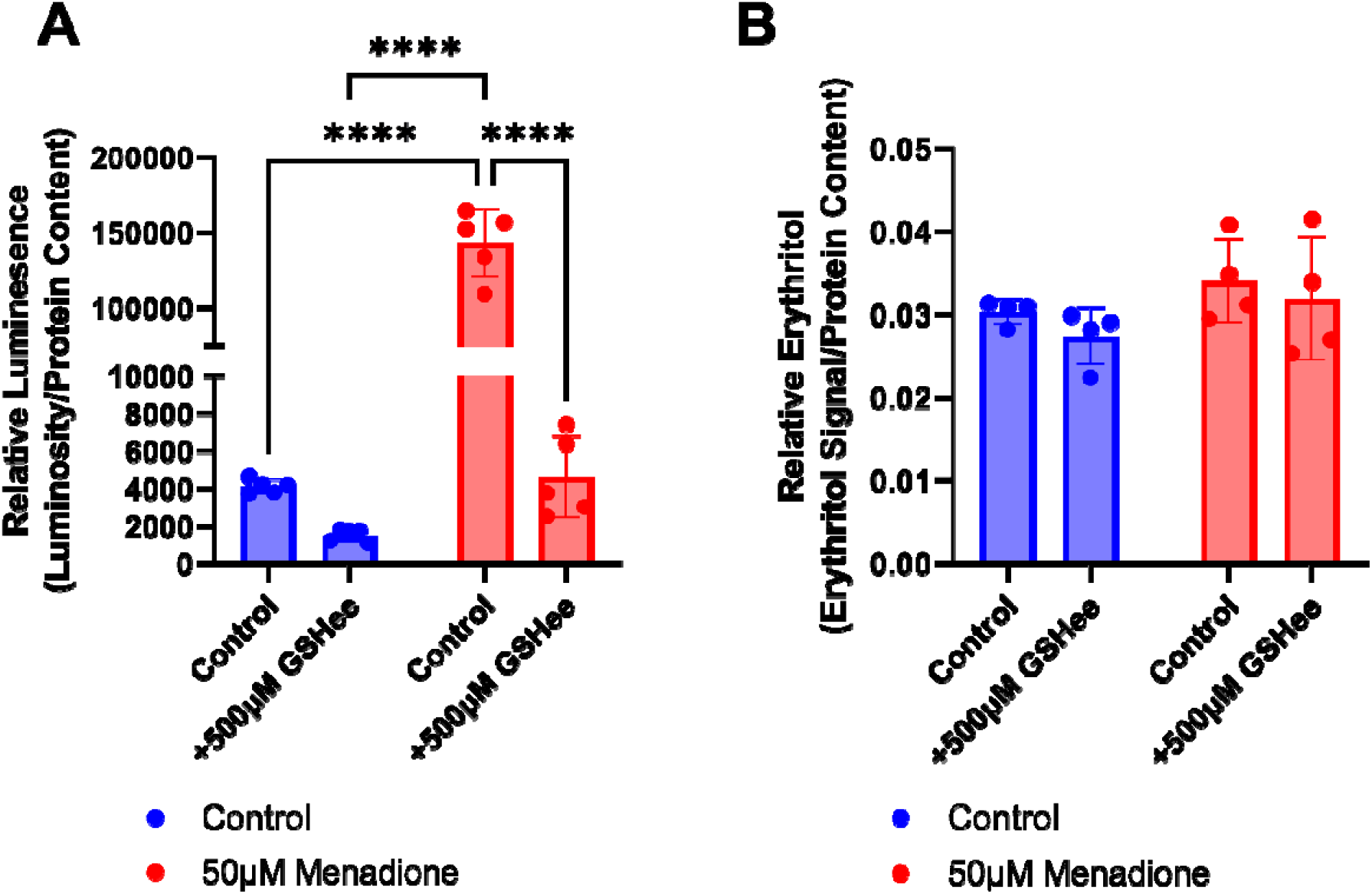
Menadione treatment does not impact intracellular erythritol in HK-2 cells. A) Relative ROS and B) relative erythritol in HK-2 cells cultured in 16 mM (high) glucose and treated with menadione and GSHee for 2 hrs. Relative metabolites were normalized to internal standard and all values were normalized to protein content. Data are shown as mean ± SD and were analyzed by two-way ANOVA followed by Tukey’s multiple comparisons test (n=4-5). ****p<0.0001. GSHee: glutathione ethyl ester; ROS: reactive oxygen species.

We have previously observed that erythritol synthesis in response to oxidative stress is not dose-dependent, but rather is maximized after a threshold for stress has been reached (3). We found that HK-2 cells were more robust when treated with ROS-inducing agents compared to C2C12 cells, suggesting a higher “threshold” for induction of erythritol synthesis despite elevated ROS. There was no elevation in erythritol synthesis with doses ranging from 50 μM to 1 mM of menadione or 1 mM of H_2_O_2_ (**Figure 5A**) in HK-2 cells, despite a dose-dependent increase in ROS with both menadione and H_2_O_2_ treatment (Figure 5B, p < 0.0001). Consistent with our observations in C2C12 cells, 1 mM H_2_O_2_ caused an elevation in intracellular sorbitol (Figure 5C, p < 0.0001), whereas menadione reduced sorbitol (Figure 5C, p < 0.05).

**Figure 5.**
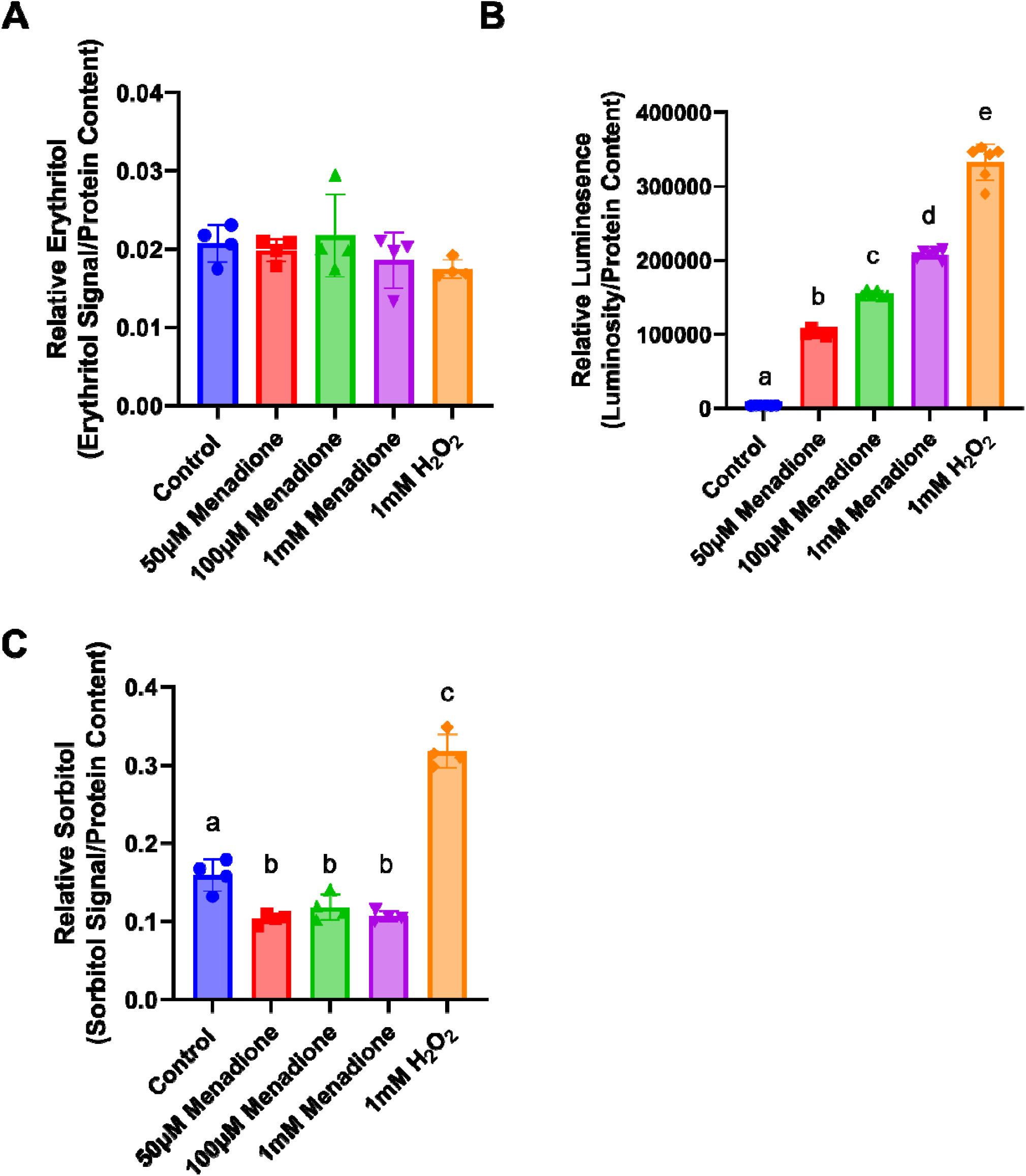
Erythritol does not respond to treatment with ROS-inducing agents in HK-2 cells. Relative A) erythritol, B) ROS, and C) sorbitol in HK-2 cells treated with menadione or H_2_O_2_ for 2 hrs. Relative metabolites were normalized to internal standard and all values were normalized to protein content. Data are shown as mean ± SD and were analyzed by two-way ANOVA followed by Tukey’s multiple comparisons test (n=4-6). Bars with dissimilar letters are significantly different (p<0.05). H_2_O_2_: hydrogen peroxide; ROS: reactive oxygen species.

### Inhibition of the polyol pathway does not increase erythritol synthesis in C2C12 or HK-2 cells

Because SORD participates in both the polyol pathway and erythritol synthesis, we hypothesized that inhibiting the polyol pathway may cause an increase in the erythritol-synthesizing activity of SORD (Figure 1). Sorbinil inhibits aldose reductase, the rate-limiting enzyme of the polyol pathway. Inhibiting aldose reductase reduces the synthesis of sorbitol, which SORD converts to fructose.

In C2C12 cells, we observed that treatment with menadione elevated erythritol, whereas treatment with H_2_O_2_ elevated sorbitol levels. We aimed to determine if inhibiting the polyol pathway with Sorbinil during H_2_O_2_ treatment could promote glucose to be redirected toward erythritol synthesis. As expected, we found that Sorbinil significantly reduced intracellular sorbitol in C2C12 cells with or without H_2_O_2_ treatment (**Figure 6A**, p < 0.0001). Rather than an increase, we observed a decrease in erythritol following Sorbinil treatment (Figure 6B, p < 0.05 in control and H_2_O_2_ treated cells).

**Figure 6.**
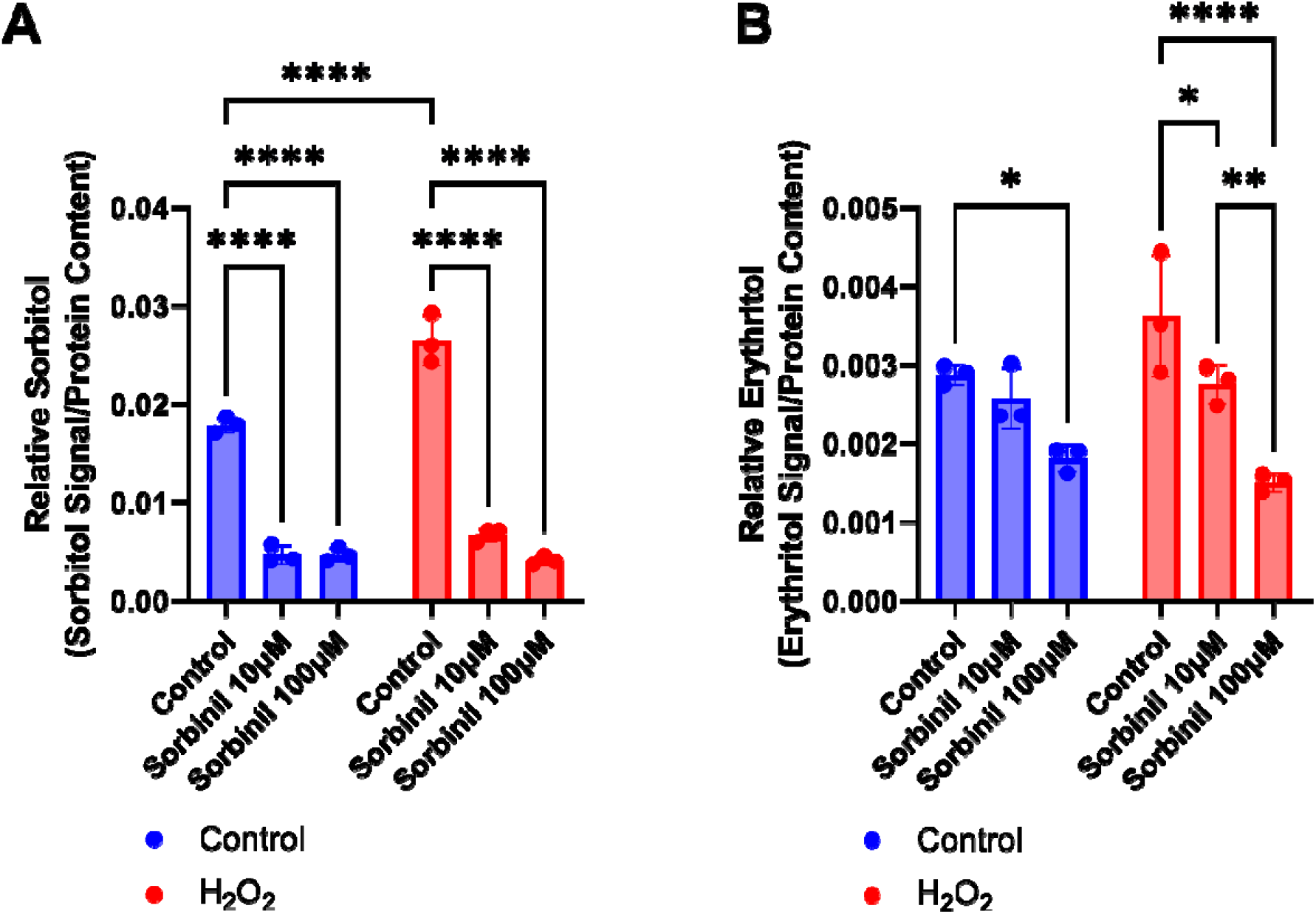
Sorbinil treatment inhibits erythritol synthesis in C2C12 cells. Relative A) sorbitol and B) erythritol in C2C12 cells following overnight treatment with Sorbinil. Relative metabolites were normalized to internal standard and protein content. Data are shown as mean ± SD and were analyzed by two-way ANOVA followed by A) Tukey’s and B) Sidak’s multiple comparisons test (n=3). *p<0.05, **p<0.01, ****p<0.0001.

We then assessed this relationship in HK-2 cells. Because stimulating oxidative stress had no impact on erythritol synthesis in HK-2 cells, we chose to perform Sorbinil treatment under low or high glucose conditions. Again, Sorbinil treatment significantly reduced intracellular sorbitol in low and high glucose (**Figure 7A**, p < 0.001 and p < 0.0001 respectively). HK-2 cells had no difference in erythritol content following Sorbinil treatment, regardless of the glucose concentration (Figure 7B).

**Figure 7.**
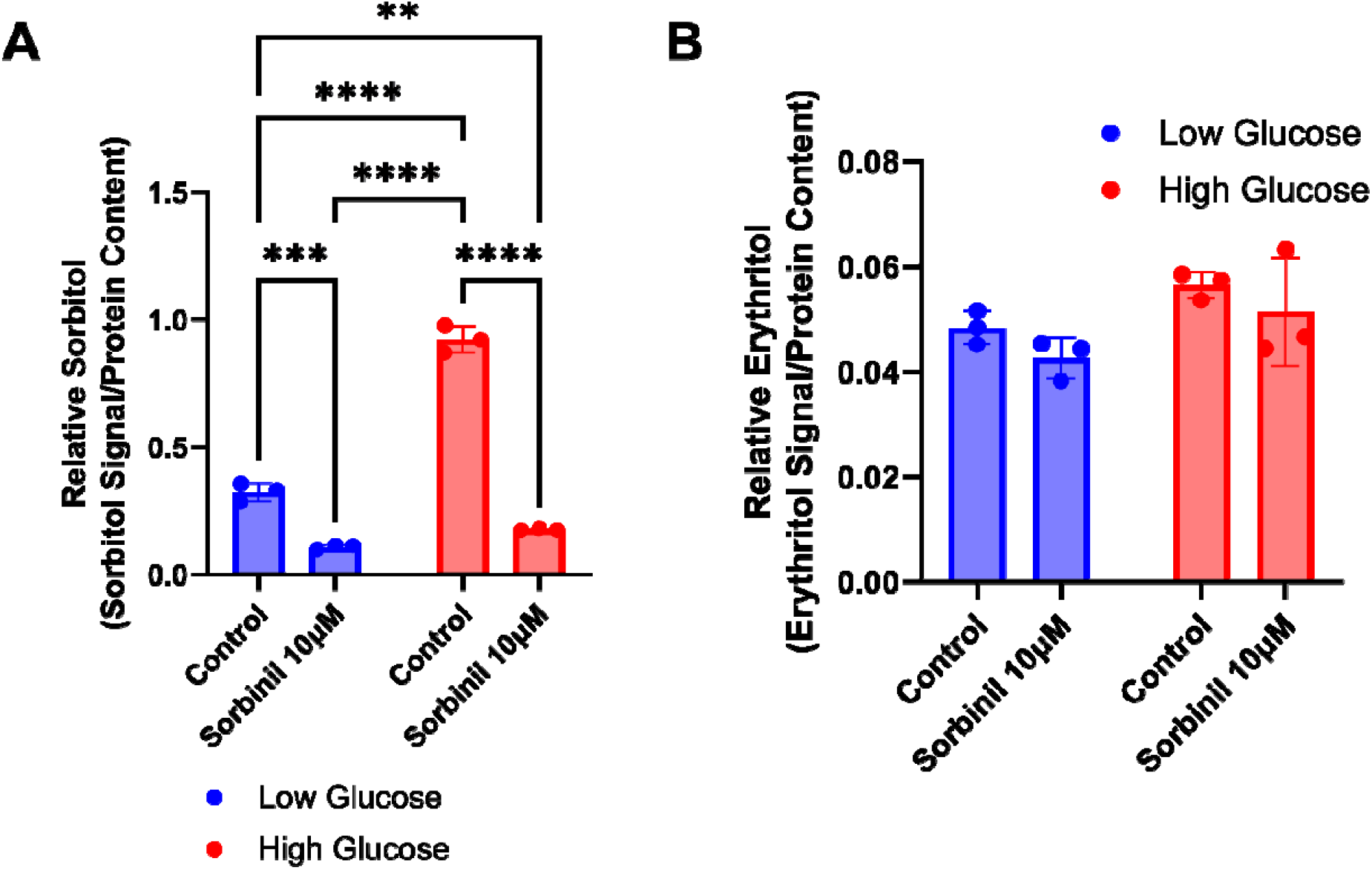
Sorbinil treatment does not impact intracellular erythritol in HK-2 cells. Relative A) sorbitol and B) erythritol following overnight treatment with Sorbinil in HK-2 cells cultured in low (5 mM) or high (16 mM) glucose. Data are shown as mean ± SD and were analyzed by two-way ANOVA followed by Tukey’s multiple comparisons test (n=3). **p<0.01, ***p<0.001, ****p<0.0001

## Discussion

The cell types which synthesize erythritol in mammals remain largely unexplored. The purpose of this study was to explore mechanisms shown to promote erythritol synthesis in cancer cells in more physiologically relevant, non-transformed kidney and skeletal muscle cells.

Consistent with our findings in A549 and HK-2 cells, we have observed that high-glucose media promotes erythritol synthesis in C2C12 cells (3). This supports that glucose availability is a key determinant of erythritol levels in mammalian cells.

Importantly, we have also observed that there are cell-specific differences in erythritol synthesis following exposure to ROS. In C2C12 cells, menadione elevates erythritol, whereas H_2_O_2_ treatment does not (Figure 3B). H_2_O_2_ treatment consistently elevated erythritol content in A549 human lung cancer cells (3). Surprisingly, HK-2 cells did not respond to ROS induction by menadione nor H_2_O_2_ treatment with elevated erythritol (Figures 4B and 5A). The PPP is an important source of NADPH to maintain reduced glutathione to combat oxidative stress (1,8,9). There are, however, many mechanisms to mitigate elevated ROS, including catalase enzymes and nonenzymatic antioxidants (10). These mechanisms, combined with baseline PPP flux, may be sufficient to mitigate ROS in HK-2 cells without additional PPP flux (and subsequent erythritol synthesis). In the skeletal muscle, baseline PPP activity is relatively low compared to other tissues (6,7). Under oxidative stress, then, a compensatory elevation in PPP flux could explain the observed increase in intracellular erythritol. These data highlight the cell-specific differences in erythritol synthesis, which is an important factor in understanding the tissue sources of high circulating erythritol.

We consistently observed that treatment with hydrogen peroxide induced elevated sorbitol levels (Figures 3B and 5C). This is likely due to the inhibition of glyceraldehyde-3-phosphate dehydrogenase (GAPDH) by hydrogen peroxide (11). GAPDH is an enzyme which separates “upper” from “lower” glycolysis. Inhibition of lower glycolysis (GAPDH) by ROS supports the buildup of glucose carbons which can be utilized through the PPP, rather than the TCA cycle (11). The observation that H_2_O_2_ stimulated sorbitol, but not erythritol, accumulation suggests that inhibiting glycolysis may not be sufficient to cause glucose “overflow” into erythritol synthesis. This observation should be validated further using stable isotope tracers and specific inhibitors of glycolytic enzymes such as GADPH or pyruvate dehydrogenase.

SORD is involved in both the polyol pathway (its primary function) and in catalyzing the conversion of erythrose to erythritol in the PPP. We have observed conditions in which the polyol pathway is elevated while erythritol remains steady, or erythritol is elevated while the polyol pathway is steady (3,5). We hypothesized that there may be a trade-off, depending on the stress applied, between SORD catalyzing fructose or erythritol synthesis. Contrary to our hypothesis, inhibition of the polyol pathway had either no effect on or reduced intracellular erythritol depending on cell type (Figures 6 and 7). This suggests that reducing sorbitol availability does not promote the erythritol-synthesizing activity of SORD. The reduction in erythritol may be due to off-target effects of Sorbinil impacting SORD activity, or downregulation of SORD in response to low aldose reductase activity. Overall, these results suggest that erythritol synthesis is not directly stimulated to cope with an abundance of glucose but that it may be caused indirectly by the accumulation of PPP intermediates.

One limitation of this work is the use of single cell types. Tissues are composed of a diverse array of cell types, which can have differing physiological roles and metabolism. C2C12 cells and similar skeletal muscle models rely heavily on anaerobic glycolysis (12). More aerobic skeletal muscle may respond differently to factors we have observed to stimulate erythritol synthesis. Additional characterization of erythritol content in red and white skeletal muscle could address this issue. Similarly, HK-2 cells represent only the proximal tubules of the kidney. There are clear differences in fuel utilization between kidney cell types (13). In vivo, proximal tubule cells preferentially oxidize fatty acids, which also may not be well represented in our *in vitro* model (13). We also did not observe a significant difference in intracellular erythritol between low and high-glucose treated HK-2 cells, which is inconsistent with our previous findings (3). This is attributed to the use of a 50-50 ratio of DMEM/F12 in the present study, which resulted in a final concentration of 16 mM rather than 25 mM glucose. Another limitation is the measurement of a single reactive oxygen species to represent oxidative stress. Reactive oxygen species are diverse molecules with distinct signaling functions and methods of detoxification. This limitation, along with the transient nature of ROS, may explain why we did not observe increased relative ROS in the C2C12 cells after menadione treatment. Although total H_2_O_2_ is a good indicator of relative oxidative stress, further exploration of erythritol and ROS would benefit from measuring additional outcomes such as the activity of antioxidant enzymes or lipid oxidation.

## Conclusion

Our findings provide further evidence that glucose availability promotes erythritol synthesis. This is a novel and particularly relevant finding in skeletal muscle cells, which are a primary site of glucose disposal. In addition, we have identified that ROS promotes erythritol synthesis in muscle myotubes, but not kidney proximal tubule cells. Finally, we have shown that there is no trade-off between polyol pathway glucose overflow and PPP erythritol synthesis. Overall, our data supports that the kidneys and skeletal muscle can contribute to erythritol production, and that the regulatory mechanisms governing erythritol synthesis vary by cell type.

## Abbreviations

DMEM: Dulbecco’s Modified Eagle Medium
H_2_O_2_: hydrogen peroxide
PPP: pentose phosphate pathway
ROS: reactive oxygen species
SORD: sorbitol dehydrogenase

## Author Contributions

SRO and MSF designed research; SRO conducted research and analyzed data; SRO and MSF wrote the paper. MSF had primary responsibility for final content. All authors have read and approved the final manuscript.

Data described in the manuscript will be made available upon request.

